# Learning Mitigates Genetic Drift

**DOI:** 10.1101/2020.06.24.170258

**Authors:** Peter Lenart, Julie Bienertová-Vašků, Luděk Berec

## Abstract

Genetic drift is a basic evolutionary principle describing random changes in allelic frequencies, with far-reaching consequences in various topics ranging from species conservation efforts to speciation. The conventional approach assumes that genetic drift has the same effect on all populations undergoing the same change in size, regardless of different behaviors and history of the populations. However, here we reason that processes leading to a systematic increase of individuals’ chances of survival, such as learning or immunological memory, can mitigate the effects of genetic drift even if the overall mortality rate in the population does not change. We further test this notion in an agent-based model monitoring allele frequencies in a population of prey, either able or not able to learn. Importantly, both these populations start with the same effective size and have the same and constant overall mortality rates. Our results demonstrate that even under these conditions, learning can mitigate drift if generations overlap. Furthermore, this effect holds regardless if the population is haploid or diploid or whether it reproduces sexually or asexually. Therefore, our findings demonstrate that learning is an overlooked factor affecting the effective population size. These findings may be of importance not only for basic evolutionary theory but also for other fields using the concept of genetic drift.

## Introduction

Genetic drift is a change in allele frequencies due to random population events. It is generally thought to depend on the population size and number of reproducing individuals, and not much else^1–3^. The key concept used to quantify the effect of drift on real-world populations is the effective population size introduced by Sewall Wright in 1930s^4,5^. While there are many extensions of the Wright’s work allowing calculation of the effective population size under different ecological scenarios ^3^, none of them considers effects of non/reproductive behaviors and history of a given population. Just as in other fields of biology^6,7^ such a simplification has proven to be very useful, but as we show further in this article, it is likely incomplete. Indeed, just as phenotype of an individual animal may change and develop during its life, so can change its response to various potentially deadly situations. This is crucial because if such a change is systematic, it can generate transient structures of different vulnerabilities to a given source of mortality in the population, and that can change how a random source of mortality affects allelic frequencies even when overall mortality rate at the population level remains the same.

Let us imagine a hypothetical scenario. Consider a species subject to predation, with all individuals genetically identical so that there is no selection acting upon them. Every young individual of this prey species has the same chance to escape a predator if it is attacked, say 50%. However, if an individual survives a predator’s attack, it learns from this experience and is more cautious or skillful in the future, increasing its chances to survive a future encounter of a similar kind (e.g., to 75%). Moreover, every future attack that the prey individual survives allows it to learn from it and further increases its chances to survive the next one (albeit at a decelerating rate). In other words, individuals which survive and learn from such attacks increase their fitness by gaining an advantage compared to those which have not yet encountered predators, even though this advantage has random origins and is not heritable. As a consequence, alleles carried by a lucky individual get a temporary boost. It is thus less likely that these alleles are removed from the population. Furthermore, because such an effect is random, all alleles benefit at some point, which, in the long run, protects them against the effects of genetic drift.

The scenario with the prey learning to escape predators is just one specific example, and the same principle can be applied to other situations and effects. For example, a population affected with a deadly pathogen in which individuals that survive infection increase their immunity should show the same trend. In this article, we test just this hypothesis that a systematic increase of organisms’ chances of survival (e.g., learning, immunological memory) in response to a random source of mortality can mitigate genetic drift even when the mortality rates for both learning and non-learning scenarios remain the same. For this purpose, we develop an agent-based model that monitors allele frequencies in time in a population of prey, either able or not able to learn how to avoid or escape predators more effectively from the experience of unsuccessful past predator attacks. Both of these populations start with the same effective size and have the same overall mortality rates.

## Methods

The agent-based model developed for this article monitors allele frequencies in time in a population of prey, either able or not able to learn how to avoid or escape predators more effectively from the experience of unsuccessful past predator attacks. It has these properties:

1. There are no mutations, migration, or selection in this model. This is intentional as we aim to study the effect of learning on the genetic drift in isolation from other evolutionary forces.
2. At the start of every simulation, there are no functional differences between individuals or alleles they carry. Alleles have no effect on the individuals carrying them.
3. At the start of the simulation, population size and every other parameter except the ability to learn are the same for learning and non-learning populations. Therefore, the initial effective population size is the same for both scenarios.
4. Overall mortality rates due to predation are constant and the same for both learning and non-learning populations. This is not affected by the fact that prey individuals in the learning population have, on average higher chance to escape predator attacks because predators repeat their attacks until a given number of prey individuals are killed. Therefore, predators may need to repeat their random attacks more times in the learning population; however, in the end, the same number of prey die each timestep in both the learning and non-learning population.
5. If not stated otherwise, generations overlap.

The precise workings of the models are as follows:

We start with a given number of prey individuals (1000) and an initial number of unique alleles (100). The individuals are assumed haploid and each obtains a randomly chosen allele. Moreover, each individual has a probability of avoiding or escaping the predator’s attack (initially 0.5). Each simulation runs for a given number of time steps (100). Predation is implicit: each time step, a given number of prey individuals are removed from the population – an individual is randomly selected, and its probability to avoid or escape predation is used to decide whether the predator’s attack is successful. If yes, the individual is removed from the population. This random selection of prey is repeated until given number of prey are removed from the population. The number of removed prey individuals per time step is always the same between the learning and non-learning scenario. If the attack is not successful and prey are learning, the probability of avoiding or escaping predation of the surviving prey individual is increased by a given proportion (0.75). In the default setting, this happens at the end of the timestep regardless if an individual prey survived one or multiple predator attacks. In an alternative setting, learning occurs after every unsuccessful attack. In addition, there is an upper bound on the probability to avoid or escape predation. Finally, the removed prey individuals are replaced by new ones: at the end of the time step, a surviving individual is randomly chosen (with repetition) and gives birth to one offspring until the population is restored to its original size (1000). Each offspring inherits the allele from its parent, and its probability of avoiding or escaping the predator’s attack is set to the initial value (0.5).

The haploid sexual and diploid sexual variations of the model differ from the above description only in the way reproduction is carried out. In the model with haploid sexually reproducing individuals, following predation, mating pairs are formed randomly and produce one offspring, its allele is randomly taken from one of its parents. In the model with diploid sexually reproducing individuals, each individual has two alleles, randomly chosen initially from the same set of 100 alleles as considered for haploid individuals. Following predation, mating pairs are again formed randomly and produce one offspring; its alleles are randomly taken from the respective pairs of alleles of its parents.

The above-described model was run using statistical software R, v. 4.0.1.

## Results

### The haploid asexual model

First, we tested the hypothesis that learning mitigates genetic drift on a simple model assuming haploid, asexually reproducing prey (Methods). Simulation results clearly show that ability to learn from experience significantly reduces the loss of genetic diversity caused by genetic drift (Fig 1) even though learning and non-learning populations in our model start with the same effective population size and have the same overall mortality rates (the same number of prey is removed from the population at each time step in any scenario).

**Figure 1:**
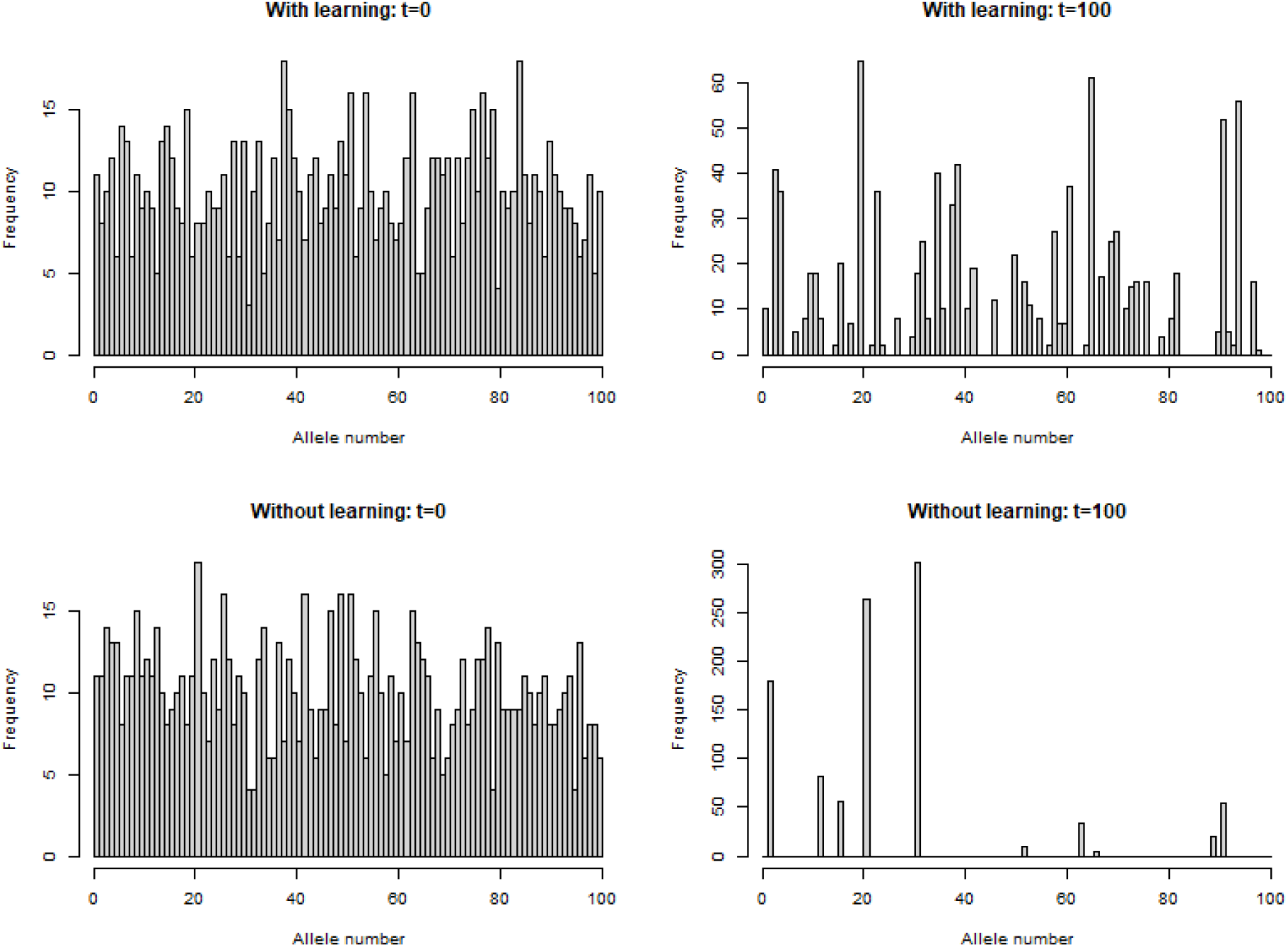
Learning slowdowns loss of genetic diversity caused by genetic drift. In this simulation run, prey populations consisted of 1000 individuals at the beginning of each step. Individuals killed by predators were replaced at the end of each step through *the* reproduction of surviving individuals (Methods). Predators consumed 500 prey individuals per timestep in both learning and non-learning scenarios to allow for a fair comparison. The upper bound on the probability to avoid or escape predator attack was set to 99%.

The size of this effect depends on specific circumstances. Our simulations consistently showed that the effect of learning on the genetic drift was affected by the fraction of the population killed by predators (Fig 2). Surprisingly, the relationship between the fraction of the prey population killed by predators and the effect of learning on the genetic drift is not always gradual (Fig 2). Instead, for any given simulation setting, there exists a narrow interval of values under which learning makes only a meager difference but above which learning greatly decreases loss of allelic diversity caused by genetic drift. Figure 2 illustrates this sudden jump by a stark contrast between simulations with 300 (Fig 2a) and 400 (Fig 2b) individuals killed per time step. The differences between simulations in which 500 (Fig 2c), 600 (Fig 2d), 700 (Fig 2e), or 800 (Fig 2f) prey individuals were killed per time step were comparatively smaller. However, as Fig 2f shows, when the intensity of predation becomes too high, learning mitigates genetic drift again less effectively. A reason for this appears to be that in such a setting, many alleles are lost in one or a few initial timesteps (initial rapid decrease in allelic diversity for both learning and non-learning scenarios, Fig 2f), which provides less opportunity for learning to affect the eventual outcome.

**Figure 2:**
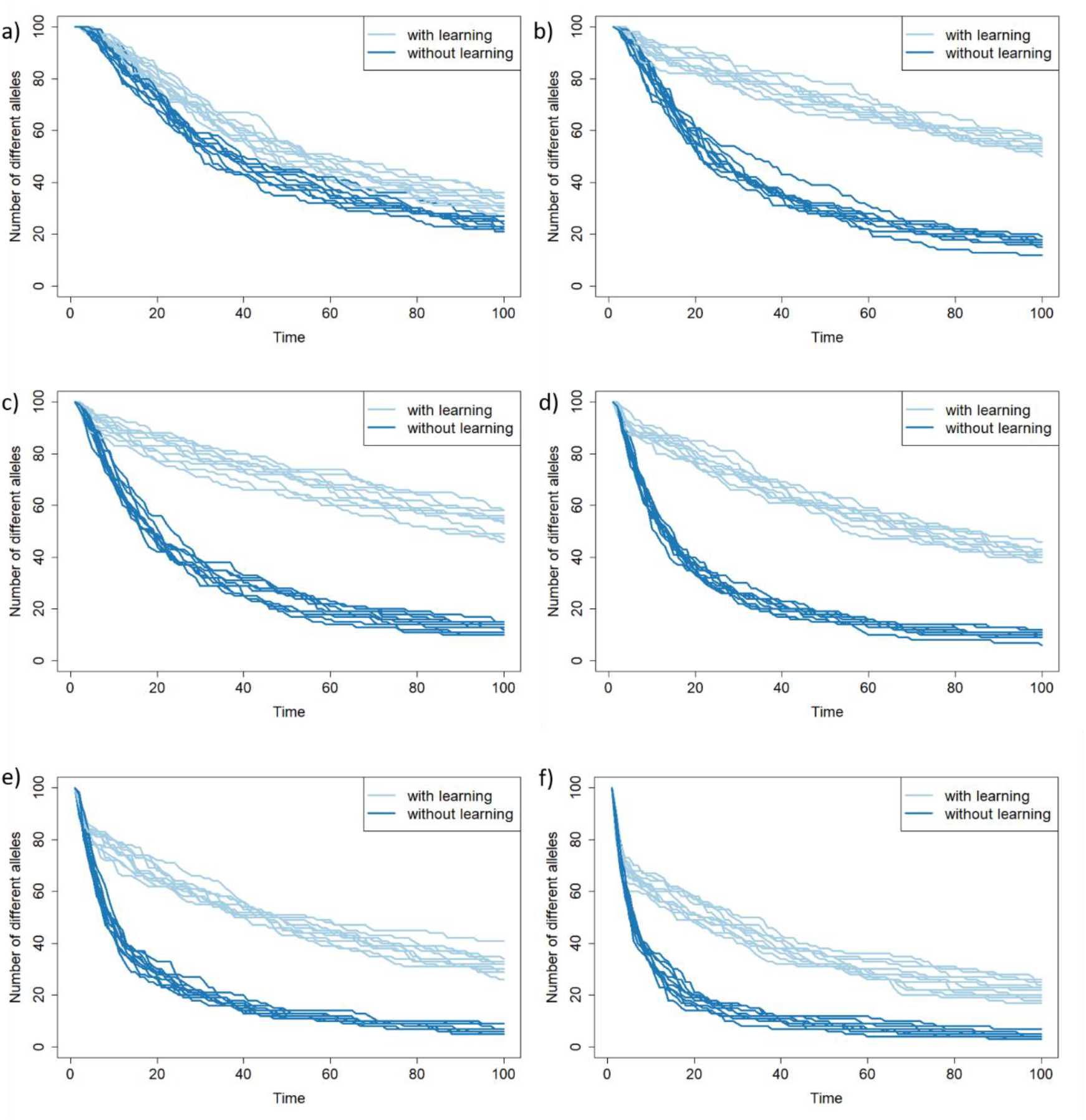
Intensity of predation modulates interaction of learning and genetic drift. Temporal dynamics of the number of alleles present in populations of prey subject to predation. All prey populations consist of 1000 individuals at the beginning of each time step. Individuals killed by predators are replaced at the end of each step through reproduction of surviving individuals (Methods). In panels a-g, predators consume 300, 400, 500, 600, 700, 800 prey individuals per time step, respectively. Predators kill the same number of individuals in both learning and non-learning scenarios to allow for a fair comparison. The upper bound on the probability to avoid or escape predator attack is set to 99%. Each panel shows ten simulation runs for both learning and non-learning scenarios.

The ability of learning to conserve genetic diversity highly depends also on how effective learning is (Fig 3). In our simulations, the maximal limit on how effectively prey could learn to avoid or escape predators was an even more important factor than the intensity of predation itself. When prey could learn at best to escape a predator attack with 90% probability, the effect of learning was noticeable (Fig 3a) but much smaller than when they could eventually reach 95% chance (Fig 3b), 97,5% chance (Fig 3c) or even 99% chance to escape (Fig 3d).

**Figure 3:**
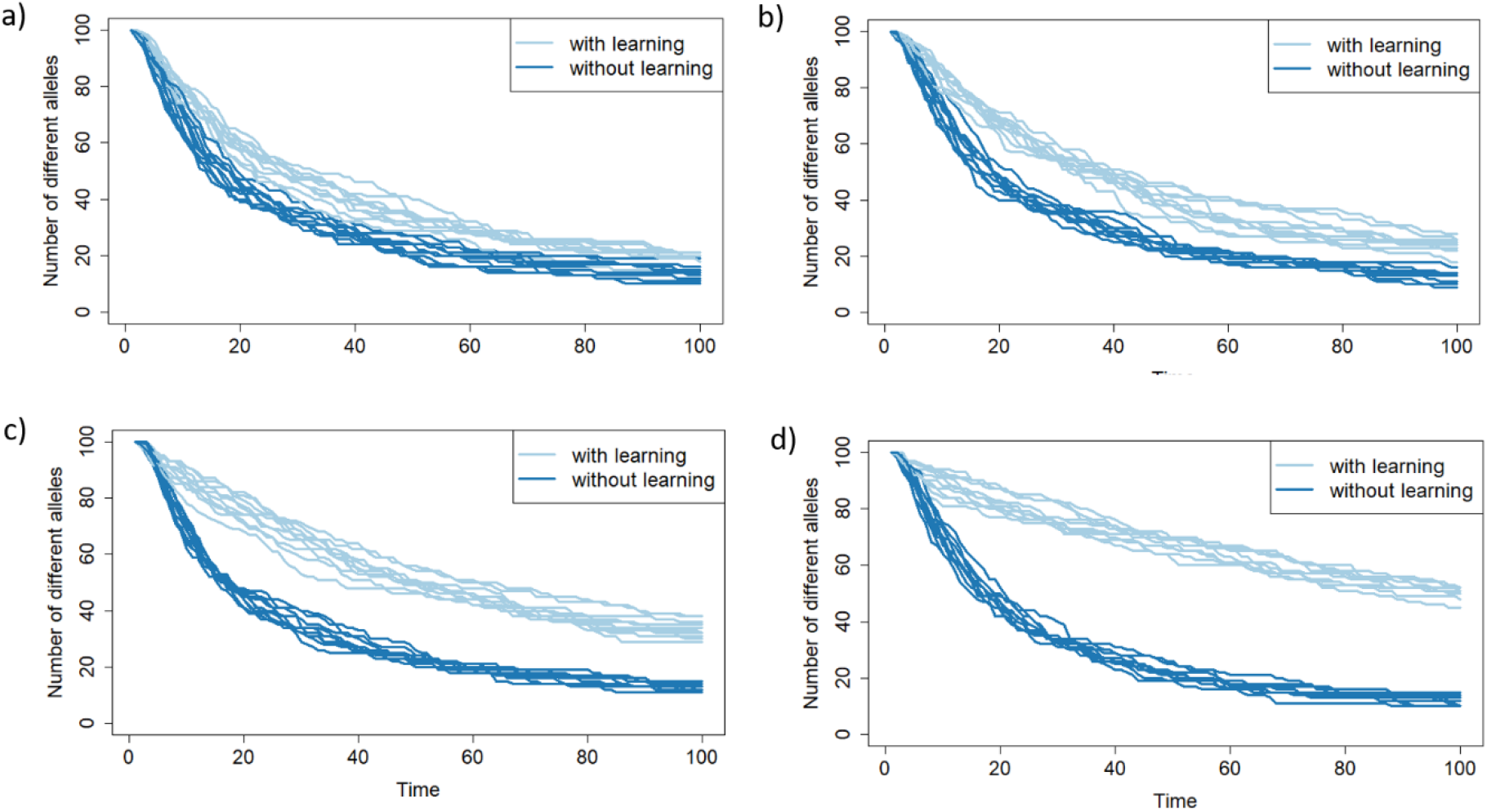
Effectiveness of learning dictates its effect on genetic drift. Temporal dynamics of the number of alleles present in populations of prey subject to predation. All prey populations consist of 1000 individuals at the beginning of each time step. Individuals killed by predators are replaced at the end of each step through reproduction of surviving individuals (Methods). Predators consume 500 prey individuals per time. Predators kill the same number of individuals in both learning and non-learning scenarios to allow for a fair comparison. The upper bound on the probability to avoid or escape predator attack is set to 90% for panel a, 95% for panel b, 97,5% for panel c, and 99% for panel d. Each panel shows ten simulation runs for both learning and non-learning scenarios.

Learning after every unsuccessful attack instead of at the end of every timestep produces qualitatively distinct results (Fig 4) that, nevertheless, supports the hypothesis that learning mitigates genetic drift. The main difference in comparison to the default setting of learning at the end of the timestep is that effect of learning is stronger when predation is less intensive (Fig 4a) and weaker when the predation is stronger (Fig 4f), i.e., opposite of the default setting.

**Figure 4:**
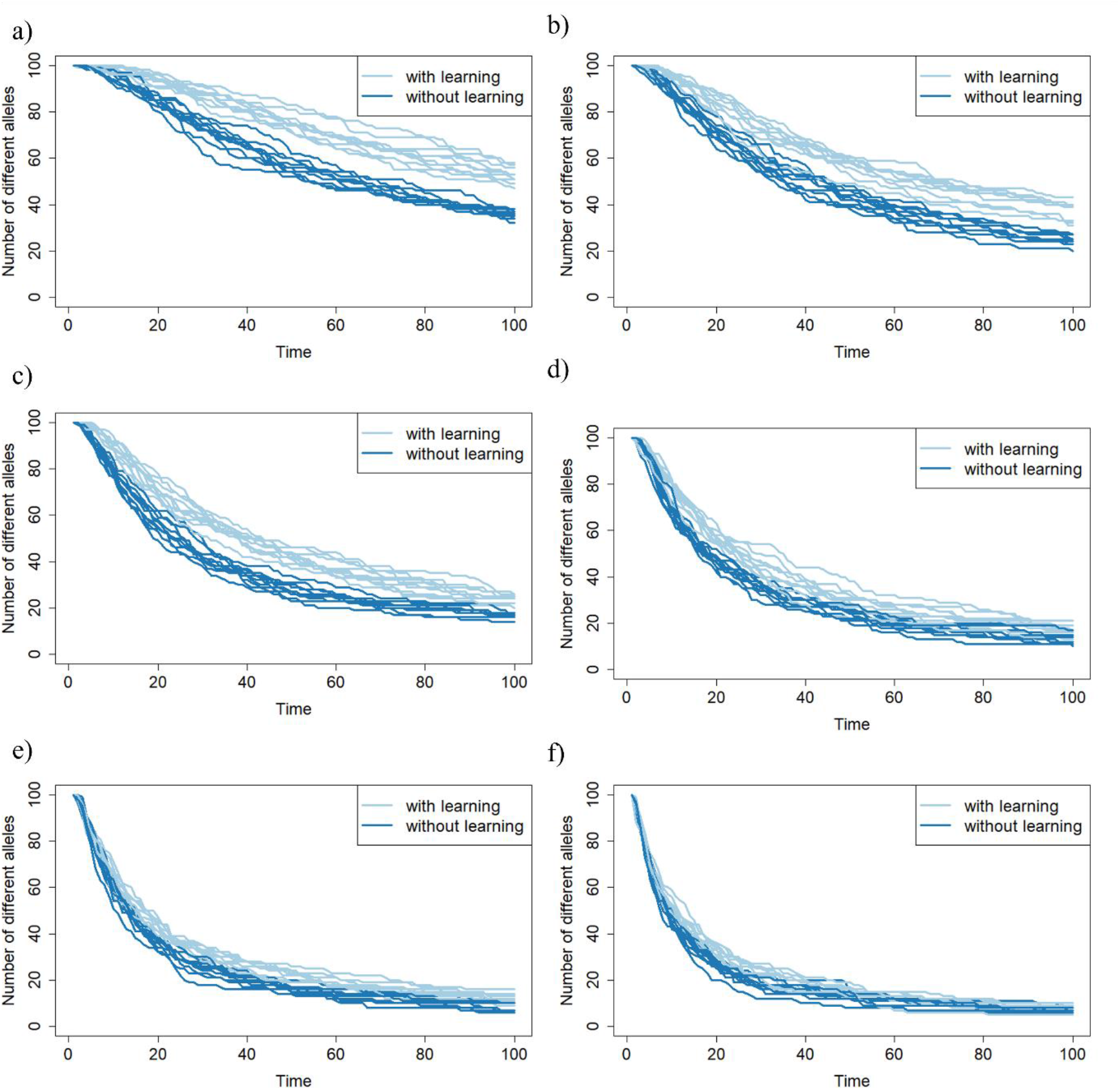
Immediate learning affects drift in a distinct way from stepwise learning. Temporal dynamics of the number of alleles present in populations of prey subject to predation. In contrast to previous simulations prey learns instantly after unsuccessful predators’ attacks. All prey populations consist of 1000 individuals at the beginning of each time step. Individuals killed by predators are replaced at the end of each step through a reproduction of surviving individuals (Methods). In panels a-g, predators consume 200, 300, 400, 500, 600, 700 prey individuals per time step, respectively. Predators kill the same number of individuals in both learning and non-learning scenarios to allow for a fair comparison. The upper bound on the probability to avoid or escape predator attack is set to 99%. Each panel shows ten simulation runs for both learning and non-learning scenarios.

At first sight, it seems likely that mean lifespan could increase in the learning scenario, which could in effect increase effective population size, thus, decreasing genetic drift. However, this is not the case, as mean lifespans in learning and non-learning scenarios are almost identical in our simulations (Table 1). In addition, the tiny difference between mean lifespan that occurs in our scenarios are most often in the opposite direction i.e., mean lifespan in learning scenario is slightly shorter.

**Table 1.** Learning scenario does not increase mean lifespan.

| no_remove | immediate_update | learning |  | non-learning |  |
| --- | --- | --- | --- | --- | --- |
|  |  | mean lifespan | SD | mean lifespans | SD |
| 200 | F | 7.258 | 13.483 | 7.304 | 13.401 |
| 500 | F | 2.812 | 9.566 | 2.960 | 8.120 |
| 800 | F | 1.828 | 6.824 | 1.860 | 6.352 |
| 200 | T | 5.056 | 7.520 | 5.252 | 4.902 |
| 500 | T | 2.956 | 8.364 | 2.978 | 8.218 |
| 800 | T | 1.871 | 6.466 | 1.865 | 6.406 |

### The haploid sexual model

To test how is the effect of learning on the strength of genetic drift affected by the mode of reproduction, we have created an updated model in which prey individuals reproduce sexually (Methods). The results from this model version were quantitatively the same as the results from the above, haploid asexual version, further supporting the notion that learning from experience is an important factor affecting the strength of genetic drift.

### The diploid sexual model

Finally, we created a version of the model with diploid sexually reproducing prey individuals (Methods). The results of this model show once again that learning from experience significantly reduces genetic drift (Fig 5). The main difference in comparison to the results of haploid model versions is that, as could be expected, genetic drift reduces genetic diversity at a slower pace.

**Figure 5:**
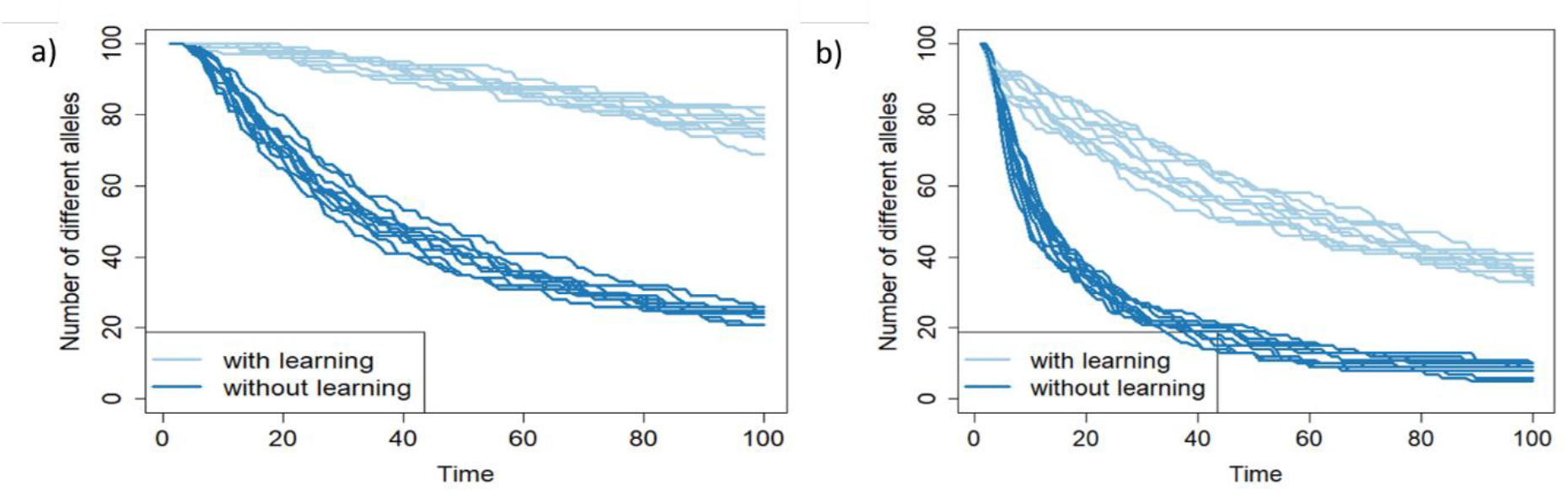
Effect of learning on genetic drift in diploid sexually reproducing population. Temporal dynamics of the number of alleles present in populations of prey subject to predation. All prey populations consist of 1000 individuals at the beginning of each time step. Individuals killed by predators are replaced at the end of each step through reproduction of surviving individuals (Methods). In panel a, predators consume 500 prey individuals per time step, while in panel b, predators consume 800 prey individuals per time step. Predators kill the same number of individuals in both learning and non-learning scenarios to allow for a fair comparison. The upper bound on the probability to avoid or escape predator attack is set to 99% for both panels. Each panel shows ten simulation runs for both learning and non-learning scenarios.

## Discussion

Genetic drift describes random fluctuations in allelic diversity in a population. In this article, we have reasoned that a systematic increase of organisms’ chances of surviving in response to a random source of mortality, e.g., learning from experience, mitigates genetic drift even when overall mortality rates remain the same. To test this hypothesis, we developed an agent-based simulation model. The results of our simulations show that even though the same number of prey die per time step in both the learning and non-learning scenarios, their allelic frequencies decrease at quite different rates. In addition to confirming our hypothesis in general, our results also demonstrate that this effect applies regardless of whether the affected organism is haploid, diploid or whether it reproduces sexually or asexually.

Our results suggest that learning from experience is an overlooked factor affecting effective population size. Contrary to most other factors such as an unequal number of males and females ^4,8,9^, the ability to learn from the experience apparently increases the effective population size as a population able to learn loses its genetic variability at a slower rate than the ideal population and thus acts as if it was a bigger ideal population. However, as the effect of learning on the genetic drift is highly dependent on many factors such as source of mortality, effectiveness of learning, etc., developing a method using information about learning to better estimate effective population sizes requires further research.

The purpose of this article is to point at the existence of a complex relationship between various mechanisms improving organisms’ ability to survive and the strength of genetic drift. The results presented here cannot, obviously, be expected to tell us anything about how strong this effect is in nature. With regard to learning from experience, the crucial factor for determining this would be assessing how effectively animals can learn to avoid or escape a deadly danger. For simulations in this article, we operated with an efficiency of 90% or even more. Arguably, this probability may be too high in real-world settings. However, under certain circumstances, it is plausible. For example, a study on barn owl has shown that these owls have a 90% chance of catching a stationary food item and only a 21% chance of catching moving items ^10^. Furthermore, the owls were even less likely to catch an item moving towards them (18%), and most interestingly, if the food items were moving sideways the barn owls were not able to catch them at all. Therefore, it seems feasible that prey may learn simple strategies that would allow them to escape at least some types of predators with surprisingly high probability. Nevertheless, even precise information about how effectively animals can learn to avoid or escape different sources of mortality may not be enough to predict the effect of learning on the genetic drift in a real-world population because the ability to learn is by itself an adaptive trait and including selection into the equation would further complicate the picture. One way or another, our message remains the same; predicting the strength of genetic drift is not as straightforward as conventionally assumed.

At first sight, it may seem that the effect of learning presented in this article is caused merely by that learning increases generational interval (the average age of parents when their offspring are born), which is known to results in increased effective population size and thus smaller genetic drift ^9^. However, as we have shown, the difference in mean lifespan between learning and non-learning scenario is not big enough to explain the observed effect. In addition, the mean lifespan in the learning scenario is, in fact, most often slightly shorter than that of the non-learning scenario – the opposite of what would be needed for the difference in lifespan to explain the weaker drift in the learning scenario.

We exemplified our idea of learning mitigating genetic drift by considering the effect of learning in the context of predation. However, for the same reasons, learning exerts the same influence on genetic drift caused by any source of mortality (or sterility), which individuals of a given species may learn to avoid. For example, if individuals of a given species were able to learn to avoid traffic from their experience, the effect on genetic drift would be identical with the learned ability to avoid or escape predators. To go even further, any learned improvement in the ability to find a partner for mating, to find food and thus decrease the risk of starvation, or to care for offspring and thus increase their chance of reaching adulthood, will, to some extent, help preserve genetic diversity despite the raw effects of drift. Accordingly, there may be other types of reactions, such as immunological memory, which could help preserve genetic diversity by mitigating genetic drift through the same principle.

The here-described relationship between learning and genetic drift also leads to some testable predictions. First, species that can learn more effectively should, on average, better conserve genetic diversity than species of less capable learners living in populations of similar size. The second and connected prediction is that species that are “better learners” should have a lower risk of extinction. This prediction is in good agreement with a recent study showing that bird species with greater behavioral plasticity indeed have a lower risk of extinction ^11^. The third prediction is that causes of excess mortality, which individuals of a given species may learn to avoid, reduce genetic diversity to a smaller degree than mortality causes which members of that species cannot learn to avoid or escape. The analogous predictions can also be made with regard to immunological memory or any process reducing mortality by a systematic reactive mechanism.

Overall, our results show that the conventional view considering genetic drift as independent of underlying species behavior is incomplete and that genetic drift may be affected by common processes such as learning or immunological memory even if overall mortality rates remain the same. Furthermore, we have shown that the level of protection against genetic drift varies in different situations, suggesting that loss of genetic diversity by genetic drift is a more complex issue than previously thought. We hope that these findings will make existing models of evolution more precise and could prove useful in a variety of topics, including the development of effective species conservation strategies, studies of the evolutionary past as well as evolutionary future.

## Acknowledgments

We want to thank Tom Kirkwood, Michael Rera, and Alan A Cohen for their valuable comments on our manuscript. The project was supported by the CETOCOEN PLUS (CZ.02.1.01/0.0/0.0/15_003/0000469) project of the Ministry of Education, Youth and Sports of the Czech Republic. The project was also supported by CETOCOEN EXCELLENCE Teaming 2 project supported by Horizon2020 (857560) and the Ministry of Education, Youth and Sports of the Czech Republic (02.1.01/0.0/0.0/18_046/0015975). As well as by the RECETOX Research Infrastructure (LM2018121).

## Code availability

The computer code used to generate results reported in this article can be accessed at https://github.com/PeterLenart/Learning_Mitigates_Genetic_Drift.

## References

1. Lequime, S., Fontaine, A., Gouilh, M. A., Moltini-Conclois, I. & Lambrechts, L. Genetic Drift, Purifying Selection and Vector Genotype Shape Dengue Virus Intra-host Genetic Diversity in Mosquitoes. PLOS Genet. 12, e1006111 (2016).

2. Lynch, M. et al. Genetic drift, selection and the evolution of the mutation rate. Nat. Rev. Genet. 17, 704–714 (2016).

3. Wang, J., Santiago, E. & Caballero, A. Prediction and estimation of effective population size. Heredity 117, 193–206 (2016).

4. Wright, S. Inbreeding and Homozygosis. Proc. Natl. Acad. Sci. U. S. A. 19, 411–420 (1933).

5. Wright, S. Evolution in Mendelian Populations. Genetics 16, 97–159 (1931).

6. Lenart, P., Scheringer, M. & Bienertova-Vasku, J. The Pathosome: A Dynamic Three-Dimensional View of Disease–Environment Interaction. BioEssays 41, 1900014 (2019).

7. Lenart, P., Scheringer, M. & Bienertová-Vašků, J. The Dynamic Pathosome: A Surrogate for Health and Disease. in Explaining Health Across the Sciences (eds. Sholl, J. & Rattan, S. I. S.) 271–288 (Springer International Publishing, 2020). doi:10.1007/978-3-030-52663-4_16.

8. Wright, S. Statistical genetics and evolution. Bull. Am. Math. Soc. 48, 223–246 (1942).

9. Hill, W. G. A note on effective population size with overlapping generations. Genetics 92, 317– 322 (1979).

10. Shifferman, E. & Eilam, D. Movement and direction of movement of a simulated prey affect the success rate in barn owl Tyto alba attack. J. Avian Biol. 35, 111–116 (2004).

11. Ducatez, S., Sol, D., Sayol, F. & Lefebvre, L. Behavioural plasticity is associated with reduced extinction risk in birds. Nat. Ecol. Evol. 4, 788–793 (2020).

